# The Lifespan Architecture of Human EEG

**DOI:** 10.64898/2026.07.28.741219

**Authors:** Milana Kasab, Sahar Allouch, Aida Ebadi, Frederic Bonnardot, Bahar Güntekin, Lütfü Hanoğlu, Görsev Yener, Ilayda Kiyi-Atilla, Veronique Paban, Birgitta Rós Ásgrímsdóttir, Paolo Gargiulo, Marc Vérin, Peter Fuhr, Ute Gschwandtner, Mohamed El Badaoui, Mohamad Diab, Mahmoud Hassan

## Abstract

Human brain maturation and aging are highly nonlinear, yet their organization at the level of large-scale electrophysiological activity remains poorly understood. We analyzed resting-state electroencephalography (EEG) recordings from 1,763 healthy individuals aged 5-85 years to map lifespan trajectories across spectral, complexity, and morphological features. Age-sensitive features clustered into distinct nonlinear trajectories, most commonly showing rapid change during childhood and adolescence followed by stabilization in adulthood, while a smaller subset exhibited turning points in midlife. These trajectories also differed in their spatial expression, ranging from highly conserved scalp-wide patterns to heterogeneous profiles in which the same feature followed distinct trajectories across scalp regions. Transition ages revealed recurring regional sequences, suggesting that diverse EEG features share common reorganization chronologies. Together, these results show that lifespan EEG variation is structured through complementary temporal trajectories, spatial architectures, and regional transition sequences. This normative framework provides a basis for investigating brain development and aging and for identifying atypical patterns associated with neurological and psychiatric disorders.

## Introduction

Across the human lifespan, the brain undergoes profound yet non-uniform structural and functional reorganization^1–3^. Development and aging reshape synaptic architecture, myelination, excitation-inhibition balance, and large-scale network communication, progressively altering the temporal dynamics through which neural activity is coordinated^4–7^. Electroencephalography (EEG), as a direct measure of population-level electrical activity, provides a window into these dynamics through complementary signal properties including oscillatory rhythms, aperiodic activity, signal complexity and waveform morphology. By capturing distinct aspects of neural coordination, these EEG features offer a unique opportunity to characterize how functional brain organization emerges, stabilizes, and declines across human life.

A broad literature has documented age-related alterations in individual EEG markers, including spectral power distributions, peak frequencies, and measures of signal complexity^8–11^. However, the global organization of typical EEG maturation and aging across the lifespan remains poorly resolved. Most existing studies have focused on restricted age ranges^9,12–15^, relied on insufficient sample sizes^9,13–18^, and have consequently contributed to restricting the documented age effects to linear shifts, despite the strongly non-linear nature of brain maturation and senescence. As a result, current knowledge remains fragmented across features, developmental stages, and analytical approaches, limiting our ability to define general principles of brain electrophysiological organization across the lifespan.

This fragmented evidence leaves several fundamental questions unanswered. Do different EEG features follow shared lifespan trajectories, or does each property age according to a distinct temporal profile? Are electrophysiological changes gradual and continuous, or do they cluster into discrete developmental and aging-related phases? When do major transitions occur across the lifespan? And are these dynamics expressed uniformly across the scalp, or do different cortical regions show distinct age-related profiles?

Here, we address these questions using a large, harmonized resting-state EEG dataset spanning childhood to late adulthood. We systematically modeled lifespan trajectories across a broad range of electrophysiological features to determine their temporal profiles, transition points, and spatial organization. By integrating these complementary dimensions, this study establishes general principles of human brain electrophysiological organization across the lifespan and provides a normative foundation for interpreting age-related variation in health and disease.

## Results

To map lifespan electrophysiological dynamics, we analyzed resting-state EEG recordings from 1,763 healthy individuals spanning 5-85 years of age. For each participant, we computed 103 spectral, complexity, and morphological features, at 19 EEG channels and as across-channel averages, enabling a high-dimensional characterization of EEG variation across development, adulthood, and aging (see Methods and Supplementary Materials Table S1 for more details about the computed features). We resolve lifespan trajectory organization using a competitive model-selection framework comparing Null, Linear, Quadratic, and Inverse models using Akaike Information Criterion (AIC)-based inference^19^. For each feature, we characterized three complementary dimensions of lifespan electrophysiological organization: (i) temporal trajectory classes describing how features evolved with age, (ii) spatial organization describing how these trajectories were expressed across the scalp, and (iii) chronological transition patterns identifying when major developmental and aging-related transitions occurred.

### Lifespan EEG dynamics follow non-linear trajectories

Across the full set of electrophysiological descriptors, age-related change was predominantly non-linear (Fig. 1). Model selection certainty, quantified using Akaike model weights (w□), showed that support was concentrated on non-linear trajectory classes (Fig. 2A). Among all feature-channel pairs, inverse trajectories, characterized by rapid early-life changes followed by stabilization, accounted for more than 40% (846/2060). Quadratic trajectories, reflecting transitions occurring later in life, were less frequently and observed in approximately 5% of feature-channel pairs. By contrast, no feature exhibited a high-certainty linear trajectory according to the Akaike weight-derived model certainty criterion, indicating that electrophysiological aging is better characterized by discrete phases rather than as a constant-rate process (Fig. 2 A).

**Figure 1.**
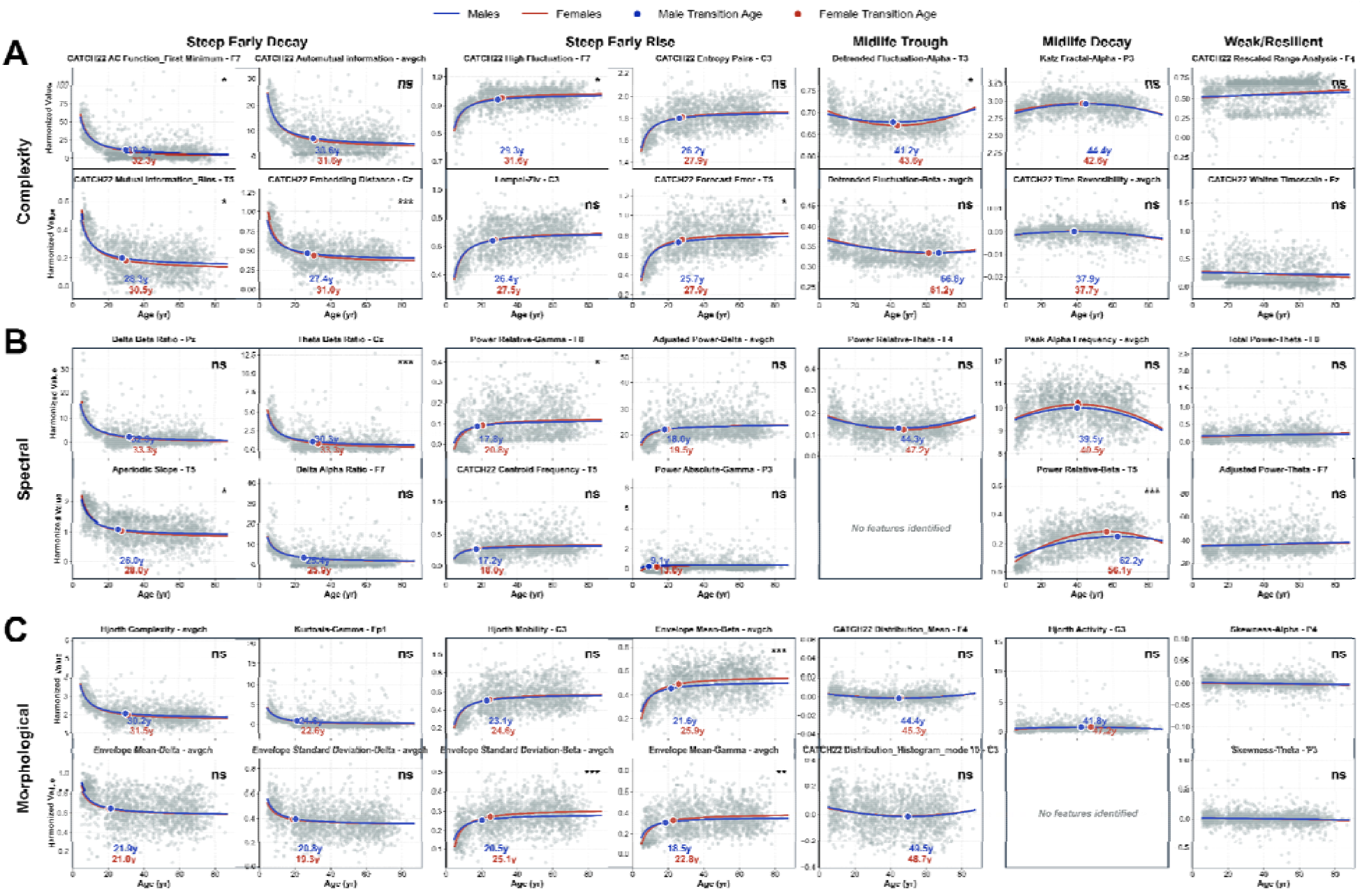
Distinct classes of lifespan electrophysiological trajectories across EEG feature domains. Representative lifespan trajectories illustrating the principal classes of age-related electrophysiological dynamics identified across complexity (A), spectral (B), and morphological (C) feature domains. Trajectories were classified into five organizational patterns: steep early decay, steep early rise, midlife restoration, midlife decay, and weak/resilient dynamics. Blue and red curves represent fitted lifespan trajectories for males and females, respectively. Colored markers indicate estimated transition or critical ages for each sex. Grey points represent individual observations. Most electrophysiological features exhibited pronounced non-linear dynamics characterized by rapid developmental transitions during childhood and adolescence followed by prolonged stabilization, whereas a smaller subset showed delayed midlife inflection points or weak age-related modulation. Statistical significance of sex effects is indicated for each feature (\**p* < 0.05, \*\**p* < 0.01, \*\*\**p* < 0.001; Benjamini–Hochberg false discovery rate (FDR) correction; ns: non-significant).

**Figure 2.**
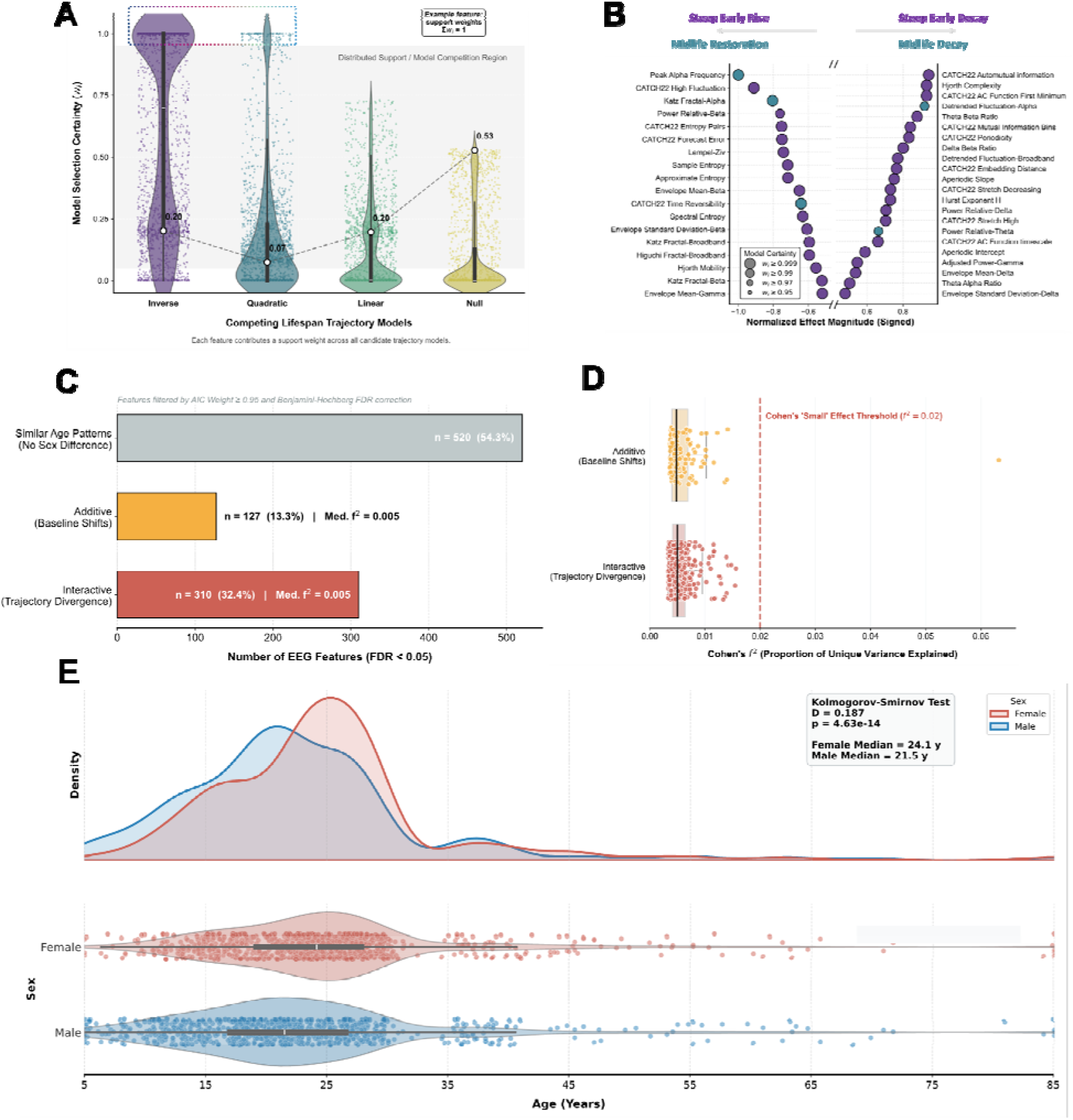
(A-B) Model selection certainty and magnitude of lifespan EEG effects. **(A)** Distribution of Akaike model weights (w⊡) across the four competing lifespan trajectory models (Inverse, purple; Quadratic, teal; Linear, green; Null, yellow). For each EEG feature, four model weights are computed—one per candidate model—representing their relative support and constrained to sum to one. Each point represents the model weight assigned to one feature–channel pair. Violin plots show the weight distributions with embedded boxplots indicating the median and interquartile range. The dashed line connects the median model weights to illustrate how model support is partitioned across the competing trajectories. The shaded grey region denotes the model competition zone, where support is distributed across multiple candidate models rather than concentrated on a single winning trajectory. The outlined region (rectangular box) highlights feature–channel pairs satisfying the high-certainty criterion (w⊡ ≥ 0.95). **(B)** Signed normalized effect magnitudes for the top 40 channel-specific features with the largest absolute effect sizes. The horizontal axis is broken to emphasize features with extreme normalized effect sizes approaching [-1.0 1.0]. Data point size is proportional to model certainty, while color indicates the winning trajectory model (Inverse, purple; Quadratic, teal). **(C-E) Assessment of sex differences in trajectory shape, baseline offset, and transition timing**. **(C)** Classification of EEG features according to sex effects following FDR correction, distinguishing shared trajectories, additive baseline shifts, and interactive trajectory differences. **(D)** Distribution of Cohen’s *f ^2^* effect sizes for the additive and interactive sex terms. Boxplots indicate the median and interquartile range; the red dashed line marks the conventional threshold for a small effect (*f ^2^*= 0.02, red dashed line). **(E)** Sex-stratified distribution of critical transition ages. Density estimates and individual observations are shown for females (red) and males (blue); the inset reports the Kolmogorov–Smirnov test.

The non-linear trajectories clustered into four main temporal classes: steep early rise, steep early decay, midlife trough, and midlife decay (Fig. 2B; Supplementary Materials Fig. S1). Each class was characterized by a distinct trajectory profile and a set of transition ages marking key periods of functional reorganization (Fig. 1). Additional feature-level trajectory plots are provided in Supplementary Materials Fig. S2.

### Steep early decay

The largest trajectory class was defined by a pronounced decrease during childhood and adolescence followed by an extended plateau in adulthood. Representative features included the First Minimum of the Autocorrelation Function, Automutual Information, Aperiodic Slope, Hjorth Complexity, and the Theta/Beta Ratio (Fig. 1A–C, “Steep Early Decay”). Most of the dynamic range of these features was lost before stabilizing at a median age of 27.6 years (range = 14.0-31.5 years). To compare effect magnitudes across features, age-related effects were expressed as normalized effect sizes (|β|; see Methods). The magnitude of the developmental decline varied considerably across features, ranging from relatively weak to highly pronounced effects (median |β| = 0.61, range = 0.12- 1.00).

### Steep early rise

A second trajectory class exhibited the opposite pattern, with rapid increases during development followed by stabilization in adulthood. Representative examples included entropy measures, Relative Beta Power, Envelope Mean-Beta, and High Fluctuation (Fig. 1A–C, “Steep Early Rise”). These features increased sharply throughout childhood and adolescence before stabilizing at a median age of 22.5 years. Developmental increases differed across features, ranging from low to highly pronounced age-related effects (median |β| = 0.51, range = 0.13-0.91). Notably, despite their opposing directions, the two early-life trajectory classes stabilized on a comparable developmental timescale.

### Mid-life decay

A third class displayed prolonged developmental trajectories extending well beyond adolescence, reaching maximal values between 29.2 and 42 years before progressively declining. Representative features included Katz Fractal dimension (alpha band) and Peak Alpha Frequency (Fig 1A–B, “Midlife Decay”). Compared with the early-life classes, these transitions occurred approximately one to two decades later. Age-related effects were predominantly high in magnitude (median |β| = 0.52, range = 0.29-1.00).

### Mid-life trough

A fourth class exhibited the inverse pattern, with features reaching minima during midlife before increasing again in later adulthood. This trajectory was observed in Detrended Fluctuation Analysis in the alpha band and Relative Theta Power (Fig. 1A–B, “Midlife trough”), which reached minima at approximately 40 and 45 years, respectively. These mid-life trough patterns exhibited a broad range of effect magnitudes (median |β| = 0.57, range=0.26- 0.95).

These four trajectory classes were not restricted to specific feature domains: spectral, complexity, and morphological descriptors were represented across all four classes. Across the full set of feature–channel combinations, 55% did not reach the high-certainty threshold for an age effect (Supplementary Table 1), including several distributional statistics, envelope-based descriptors, absolute spectral power in the delta and theta bands, and several low-frequency fractal dimensions. Importantly, age-related effects remained stable after age-bin matching stratified by sex, indicating that the observed trajectories were not driven by demographic imbalance (See Supplementary Fig. S3).

Overall, the observed patterns indicate that age-sensitive EEG features are organized into distinct nonlinear lifespan trajectories, with most characterized by rapid developmental remodeling before adulthood and a smaller subset showing turning points in midlife.

### Sex differences are expressed in transition timing rather than trajectory shape

Lifespan EEG trajectories were highly conserved across sexes (Fig. 2C). More than half of all analyzed features (54.3%) exhibited identical age-related trajectories in males and females, with no detectable sex effect (*FDR corrected*) (Fig. 2C). Among the remaining features, sex-related differences were primarily expressed as differences in the shape of the age-related trajectory (32.4% of features), as observed for Relative Beta Power and Beta Envelope Standard Deviation, whereas a smaller subset (13.3%) exhibited stable baseline offsets across the lifespan, most notably Peak Alpha Frequency. Despite these statistically significant effects, their magnitude remained uniformly small (Fig. 2D). The incremental variance explained by sex-related terms, quantified using Cohen’s f², was low for both baseline shifts and trajectory divergence (median f² = 0.005), well below the conventional threshold for a small effect (f² = 0.02)^20^. Thus, at the level of trajectory shape and baseline, males and females followed largely similar maturational and aging trajectories, with only subtle sex-dependent differences.

In contrast, sex-related differences were more apparent in the timing of electrophysiological transitions. Transition ages, derived from model stabilization and inflection points, followed a similar two-stage pattern in both sexes: a major wave during adolescence and early adulthood, followed by a smaller secondary wave in midlife (Fig. 2E). However, the distributions of transition ages differed significantly between males and females (Kolmogorov-Smirnov K-S Test, *D = 0.187, p <0.001*). Females exhibited a more synchronized and temporally constrained developmental window, with transition densities peaking sharply between 20 and 30 years of age. Males, by contrast, showed a broader and more prolonged maturation period, with peak transition density occurring approximately three years earlier and extending into the late twenties. A weaker secondary wave of transitions associated with delayed quadratic trajectories was observed in both sexes during midlife.

Overall, sex-related variation in lifespan EEG dynamics is expressed primarily through differences in the timing and synchronization of electrophysiological transitions rather than through differences in the overall shape or magnitude of developmental and aging trajectories.

### Lifespan EEG dynamics are organized into three spatial modes

We next examined how lifespan trajectories were distributed across scalp locations. Three distinct modes of spatial organization emerged, ranging from highly conserved scalp-wide dynamics to strongly heterogeneous and spatially restricted lifespan patterns (Fig. 3A; Supplementary Materials Fig. S4).

**Figure 3.**
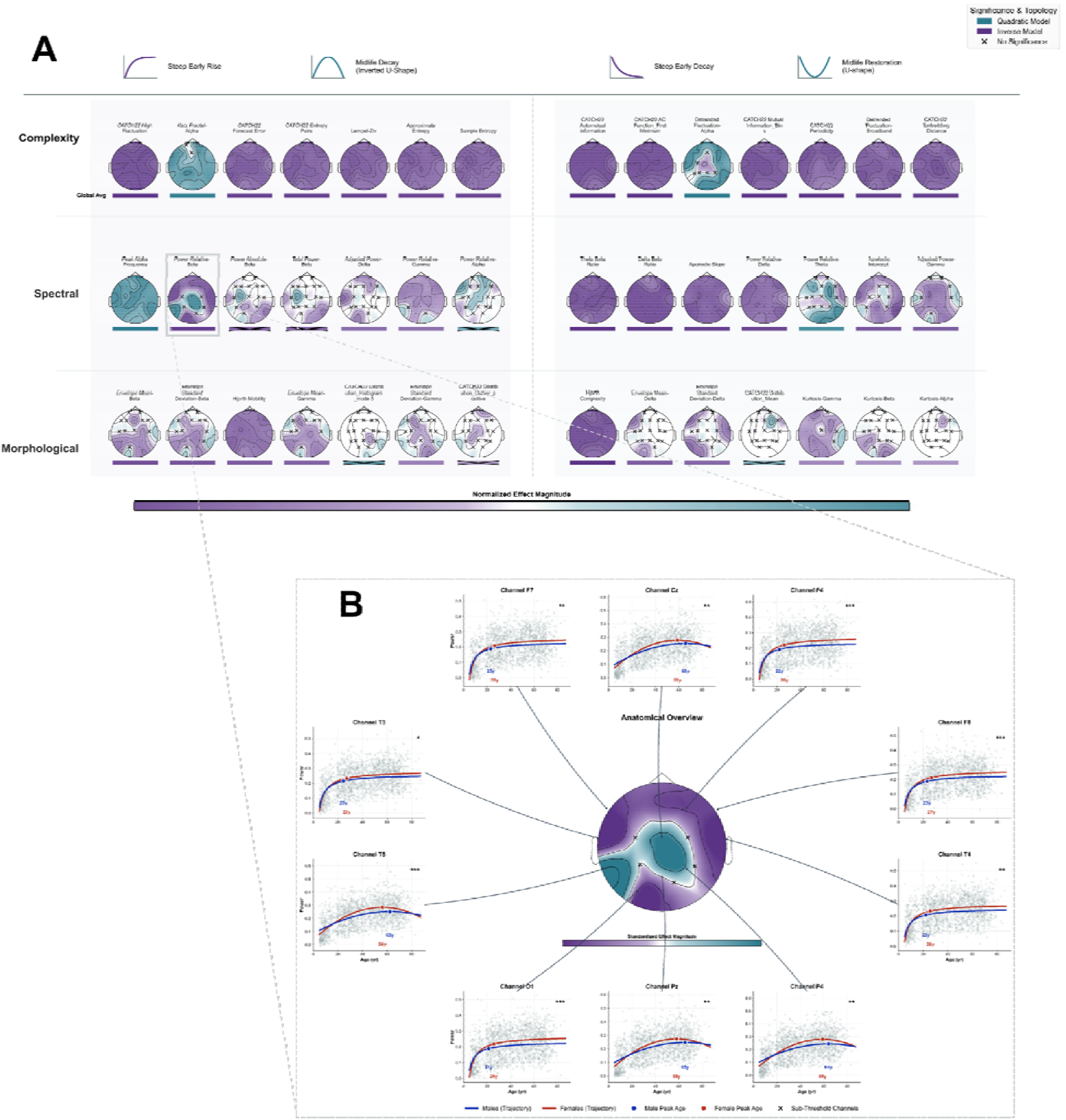
Multiscale topography of age-related changes in EEG features. **(A)** Spatial distribution of age-related effects across EEG features exhibiting high model certainty (*w*□ ≥ 0.95). Features are organized according to their dominant lifespan trajectory class: steep early rise, midlife trough (U-shape), steep early decay, and midlife decline (inverted U-shape). Each topographic map depicts the regional distribution of normalized effect magnitudes, while the horizontal bar beneath each map summarizes the corresponding whole-brain average effect. Channels not meeting the certainty threshold are marked with × symbols. Color intensity reflects normalized effect magnitude, with darker colors indicating stronger age-related effects. (B) Representative example illustrating regional heterogeneity in lifespan transitions for a single EEG feature. The central topographic map displays the spatial distribution of normalized effect magnitude across scalp locations, while surrounding channel-specific trajectory plots show fitted age-related curves for selected electrodes. Annotated ages indicate the estimated channel-level transition points, demonstrating that different scalp regions undergo electrophysiological reorganization at distinct stages of the lifespan despite sharing a common global trajectory class.

### Globally coordinated trajectories

Approximately 30% of analyzed features exhibited highly conserved developmental trajectories across the scalp. For these features, all channels converged on the same trajectory class (19/19 channels), demonstrating complete agreement between channel-level and scalp-averaged lifespan profiles (Fig. 3 A). This organization was particularly prominent among several high-effect features spanning both complexity and spectral domains, including Automutual Information, entropy measures, broadband fractal properties (Detrended Fluctuation Analysis (DFA) Coefficient, Higuchi and Katz dimensions), Peak Alpha Frequency, Aperiodic Slope, and the Theta/Beta Ratio. A complete list of features exhibiting this organization is provided in Supplementary Table 2.

This spatial consistency extended beyond trajectory shape to effect magnitude. Channel-level effect sizes exhibited minimal spatial dispersion (median dispersion = 0.07; Supplementary Table 2), indicating highly similar age-related associations across channels. Consistent with this observation, Bland–Altman analysis^21^ (see Methods) revealed strong agreement between maximal channel-level and scalp-averaged effect sizes (mean bias = 0.05; Fig. 4B), with only a small number of features, including the Delta/Beta and Delta/Alpha ratios, showing modest deviations from the limits of agreement. The strong cross-channel agreement in both trajectory shape and effect magnitude suggests that these features are dominated by globally coordinated processes, such that comparable signals could be detected across individual channels as well as in the scalp average with minimal loss of temporal or effect-size information following spatial averaging.

**Figure 4.**
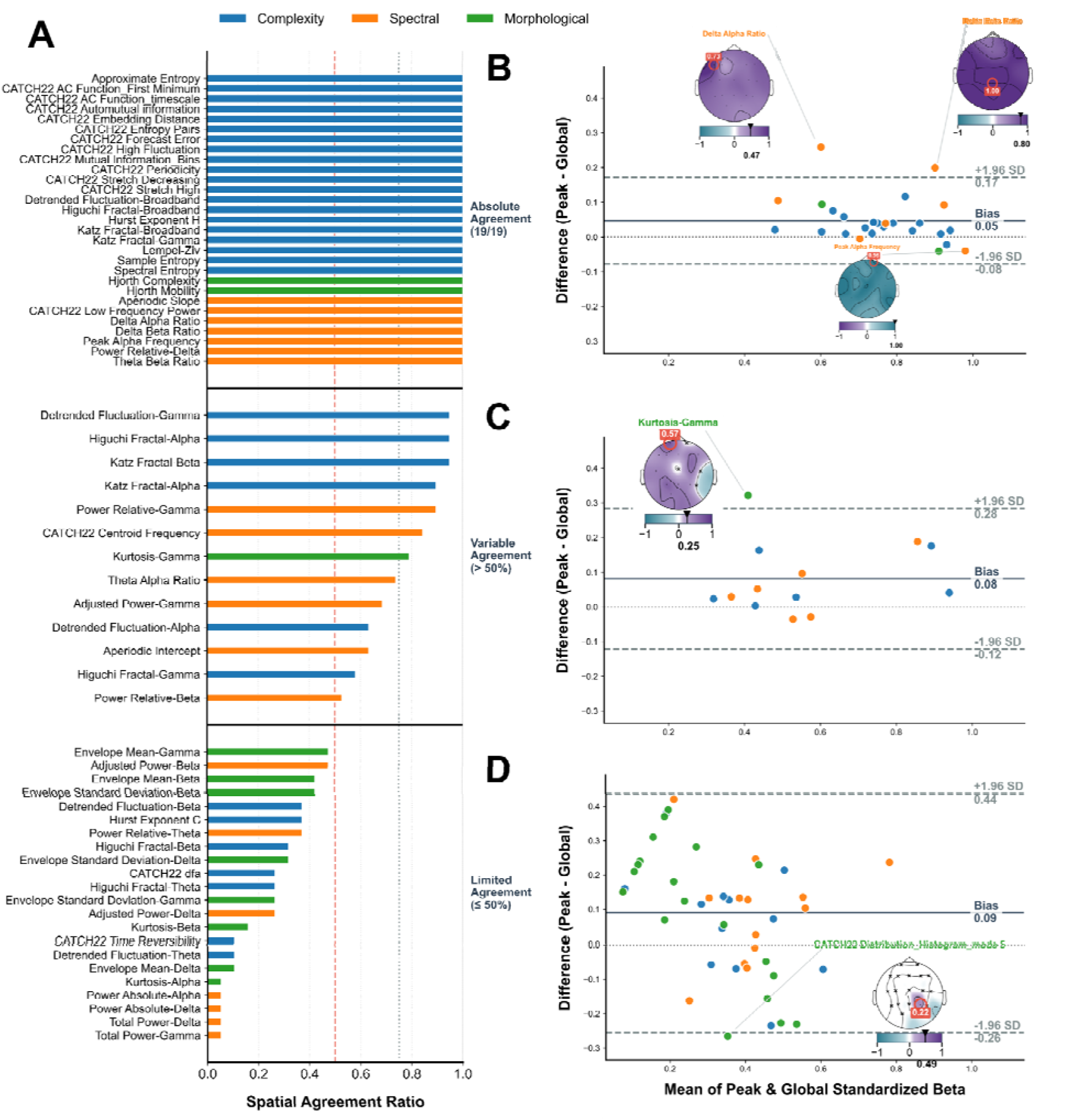
Spatial concordance between local channel trajectories and global feature-level aging dynamics. **(A)** Spatial concordance ratio for all electrophysiological features, quantified as the proportion of scalp channels (out of 19) whose locally derived lifespan trajectory matched the dominant global trajectory identified from the whole-brain average signal. Features are ranked according to the degree of local–global agreement. Colors indicate feature domains (Complexity, blue; Spectral, orange; Morphological, green). Horizontal reference lines separate features exhibiting absolute agreement (19/19 channels), variable agreement (>50% of channels), and limited agreement (≤50% of channels). Dashed vertical lines denote the 50% agreement threshold and the 75% strong-consensus threshold. **(B–D)** Bland-Altman-style assessment of agreement between global and regional age-related effect magnitudes stratified by spatial concordance level. The x-axis represents the mean normalized effect magnitude computed from the global and peak regional estimates, while the y-axis represents their difference (Peak − Global). Solid horizontal lines indicate the mean bias and dashed lines denote the 95% limits of agreement (±1.96 SD). Point colors correspond to feature domains. Representative topographic maps illustrate features within each agreement category. **(B)** High consensus features (≥19 channels). Features exhibiting near-complete spatial agreement between local and global trajectory classifications show minimal bias and narrow limits of agreement, indicating that regional age-related dynamics closely mirror whole-brain aging patterns. **(C)** Variable consensus features (10–18 channels). Features with partial regional agreement display moderate divergence between local and global estimates, suggesting increasing spatial heterogeneity in the expression of age-related effects. **(D)** Limited consensus features (<10 channels). Features with weak local–global agreement demonstrate substantial regional variability, where localized age-related dynamics deviate markedly from the dominant whole-brain trajectory. Representative examples highlight spatially restricted patterns that are not adequately captured by global averages alone.

### Spatially heterogeneous trajectories

A second mode, observed in approximately 12% of features, was characterized by the coexistence of distinct trajectory families across scalp locations. Representative examples included DFA in the alpha band, Aperiodic Intercept, and Adjusted Gamma Power, for which inverse steep-decay trajectories coexisted with quadratic U-shaped profiles across different channels (Fig. 3A). Similar spatial dissociations were observed for Relative Beta and Gamma Power, where steep early-life increases coexisted with quadratic inverted-U trajectories (Fig. 3B).

Compared with globally coordinated features, these profiles exhibited greater spatial variability in both trajectory shape and effect magnitude (median dispersion = 0.12; Supplementary Table 2). Consequently, scalp averaging captured only one component of a broader spatial organization, while alternative lifespan trajectories remained evident at the channel level. This reduced spatial concordance was reflected by channel-to-global agreement ratios ranging from 10/19 to 18/19 channels (Fig. 4A). Bland–Altman analysis^21^ further revealed a systematic tendency for channel-level effects to exceed those observed in the averaged signal (mean bias = 0.08; Fig. 4C). Although most features remained within the limits of agreement, Gamma-band Kurtosis exhibited a pronounced localized age effect that was only partially preserved after spatial averaging. Notably, these heterogeneous organizations did not follow a common topographical pattern, and distinct trajectory families were not consistently associated with specific scalp regions. These observations indicate that the apparent lifespan organization of several electrophysiological features depends critically on the spatial scale of analysis.

### Spatially localized trajectories

The most common mode, accounting for approximately 58% of the analyzed feature space, consisted of features with weak or spatially restricted age-related effects. In these cases, significant lifespan trajectories coexisted with channels that failed to reach significance, resulting in incomplete spatial concordance and reduced agreement with the averaged signal (Fig. 5A; Supplementary Table 2). This pattern was observed predominantly among morphological and spectral features, including Gamma-band Envelope Mean, Absolute Beta Power, and Alpha-band Kurtosis, where significant age effects were confined to localized scalp regions while neighboring electrodes remained non-significant (Fig. 3A).

**Figure 5.**
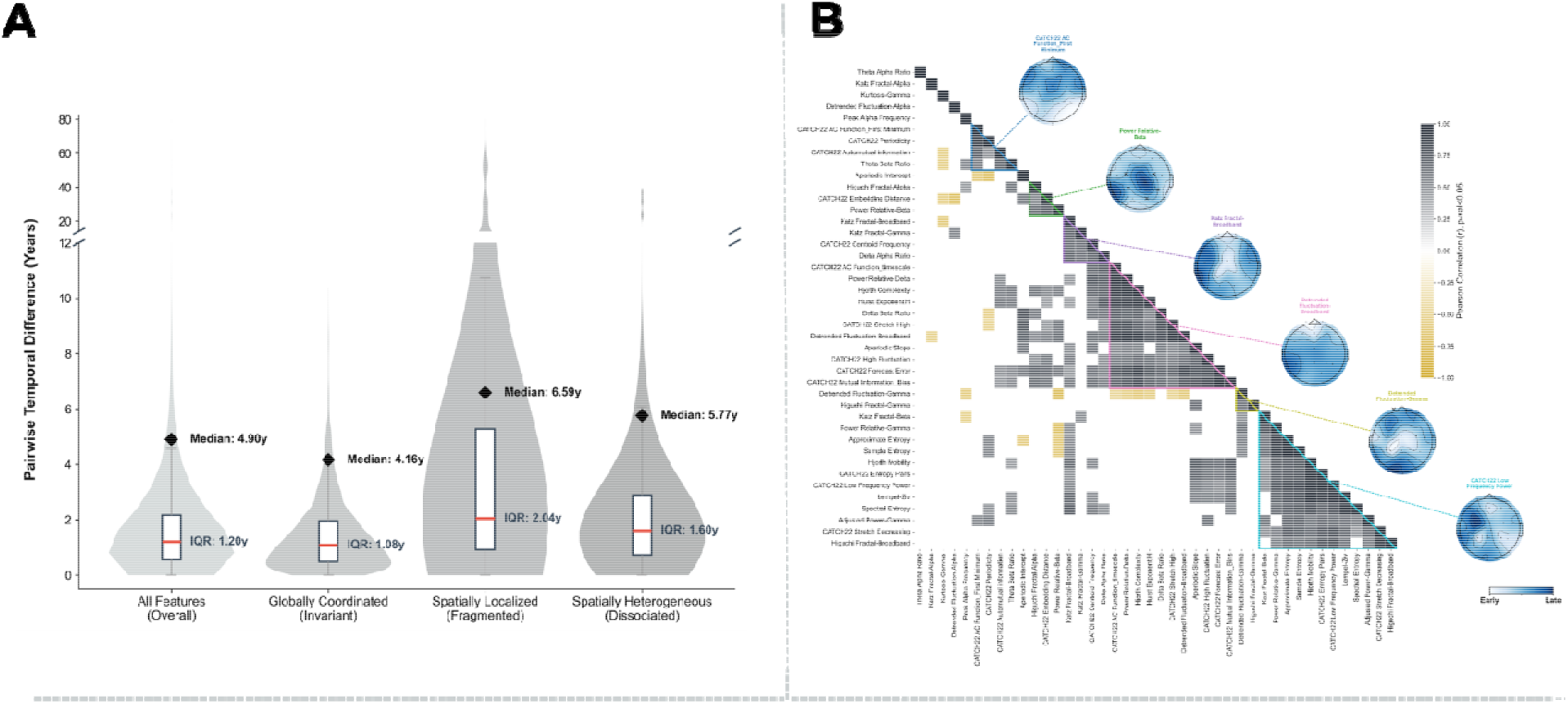
Spatiotemporal dispersion and structural clustering of EEG lifespan transitions. **(A)** Distribution of pairwise transition-age differences (in years) across scalp locations, categorized by spatial organizational mode (Globally coordinated, heterogeneous and spatially localized modes). The y-axis features a scale break to preserve the resolution of dense inner quartiles while accommodating extreme spatial transition spans. For each distribution, black diamonds represent the global lifespan sweep **(the median temporal gap between the absolute earliest and latest transitioning channels)**, while the red lines within the boxplots indicate the typical inter-regional gap (Interquartile Range, IQR). Boxplot whiskers extend to 1.5× IQR. **(B)** Lower-triangle heatmap of pairwise Pearson correlation coefficients between spatial transition maps (*p<0.05*). Features are ordered via hierarchical clustering (average linkage), revealing distinct, internally coherent co-aging spatial manifolds (highlighted by colored triangular outlines). Representative topographic maps for major clusters are projected along the main diagonal, illustrating the characteristic geometry of regional transitions progressing from early (light blue/white) to late (dark blue) developmental milestones. Channels marked with ’x’ denote mathematically sub-threshold or non-significant locations for the exemplar feature.

Correspondingly, effect sizes were generally small (median |β| = 0.14; range = 0.019–0.50) and showed the greatest divergence between channel-level and scalp-averaged estimates (Bland-Altman bias = 0.09; Fig. 4D). Depending on the spatial distribution of significant electrodes, the averaged signal could either reflect one of the channel-level trajectories (e.g., Beta-band Envelope Mean) or fail to exhibit a significant lifespan trajectory altogether (e.g., Catch22^22^ Distribution Mean). As with the dissociated profiles, these localized effects did not exhibit a consistent topographical organization across features.

### Transition timings follow conserved spatial sequences

Mapping critical transition ages across scalp locations revealed that developmental transitions did not occur simultaneously throughout the brain. In the full feature space, the median difference between the earliest and latest channel-level transition ages was 4.9 years, indicating substantial scalp-wide variability in the timing of electrophysiological reorganization. This temporal span varied by three spatial organizational modes: it was largest for spatially localized features (6.59 years), intermediate for spatially heterogeneous features (5.77 years), and smallest for globally coordinated organization (4.16 years) (Fig. 5 A).

To quantify more typical temporal offsets between regions, we examined the distribution of transition-age differences across all channel pairs-instead of considering only earliest and latest transitioning channels. Across the full feature space, the interquartile range of pairwise transition-age differences was 1.20 years, indicating that most regional transitions occurred within a relatively narrow temporal window despite the larger scalp-wide span. Pairwise dispersion was greatest for spatially localized features (IQR = 2.04 years), intermediate for spatially heterogeneous features (IQR = 1.60 years), and smallest for globally coordinated features (IQR = 1.08 years) (Fig. 5 A).

To identify regions undergoing nearly synchronous electrophysiological reorganization, we examined channel pairs whose transition-age differences fell within the interquartile range (IQR) of the pairwise transition-age distribution, thereby capturing the most closely timed regional transitions. Across the full feature space, these closely timed transitions were concentrated predominantly within frontal occipital and parietal regions. This overall organization, however, differed substantially across the three spatial modes. For globally coordinated features, the smallest differences in transition age were mainly observed between posterior channels. Spatially heterogeneous features showed closely timed transitions across a broader set of regions, particularly within frontal and central areas and between frontal, central, and occipital channels. For spatially localized features, closely timed transitions were restricted mainly to a small group of posterior channels, in line with their more limited spatial distribution.

To determine whether different EEG features shared common spatial organizations of transition timing, we clustered scalp maps of transition ages after normalizing each feature to its own developmental timeline. This approach grouped features according to the relative pattern of early and late transitions across the scalp, rather than their absolute transition ages. The analysis identified six recurring spatial profiles of electrophysiological aging (Fig. 5B).

Each profile exhibited a distinct and reproducible spatial organization of transition timing. One profile, exemplified by DFA and Higuchi Fractal (gamma band), was characterized by early transitions over central regions surrounded by progressively later transitions along the anterior-posterior axis. A second profile, represented by Katz Fractal Dimension (broadband), showed a continuous band of early transitions extending from the mid-frontal to occipital scalp. A third profile, exemplified by Relative Beta Power, and Higuchi Fractal Dimension (alpha), displayed a clear followed a peripheral-to-central gradient, with peripheral scalp regions transitioning earlier than central regions. One of the largest profiles, represented by DFA (broadband) exhibiting a pronounced anterior posterior gradient, with the latest transitions concentrated over the left parietal region. Representative maps for each profile are shown in Fig. 5B, while the complete set of feature-level transition maps is provided in Supplementary Materials Fig. S5, S6. Together, these findings indicate that lifespan EEG dynamics are organized around a limited number of conserved spatial transition profiles. Individual features differ in when transitions occur, but many follow the same spatial sequence of electrophysiological reorganization across the scalp.

## Discussion

The present findings reveal reproducible principles of human EEG maturation and aging. EEG features followed a limited set of non-linear lifespan trajectories, expressed through distinct spatial organizations and conserved transition-timing profiles. Lifespan EEG therefore appears to be structured across complementary temporal, spatial, and chronological dimensions, providing a unified framework for understanding functional brain maturation and aging.

The predominance of nonlinear trajectories is consistent with the known biology of human brain development and aging. Rather than progressing at a constant rate, brain maturation unfolds through successive periods of biological reorganization, including early synaptogenesis, extensive synaptic pruning during childhood and adolescence, prolonged myelination into early adulthood, and later cortical thinning during aging, each occurring over distinct developmental windows^4,23,24^. This nonlinear organization has been consistently observed across multiple neuroimaging modalities, with structural MRI revealing age-dependent trajectories of cortical morphology and white matter^1,24,25^, while functional MRI studies similarly report age-dependent reorganization of large-scale brain networks^3,26^. Recent MEG studies report similarly nonlinear maturation of neural dynamics across the lifespan^27^, particularly observed in the Relative Power of the Delta, Theta, Beta and Gamma Bands^27^ (Fig 1, Supplementary Materials Fig S2). Within this biological context, the apparent stabilization of many EEG descriptors during adulthood should not necessarily be interpreted as complete biological stability. Structural and functional brain organization continues to evolve throughout adulthood and aging^1,3,24^, although these changes may not be equally reflected across all EEG descriptors.

One of the most key observations was that mathematically distinct EEG descriptors repeatedly converged onto the same developmental trajectories. Spectral measures, complexity metrics, morphological descriptors, and nonlinear time-domain EEG features, including Catch22-derived measures^22^, frequently resolve into identical lifespan programs despite quantifying fundamentally different properties of neural activity. For example, descriptors capturing oscillatory dynamics: relative power of delta, theta/beta ratio (Fig. 1B), non-oscillatory dynamics: aperiodic slope & intercept (Fig. 1B, Supplementary Materials Fig S2), signal complexity: Automutual Information, periodicity, broadband DFA (Fig. 1A), and temporal structure: delta envelope standard deviation & mean, Hjorth complexity (Fig. 1C) often followed the same steep early developmental trajectories, reflecting coordinated neurobiological transitions rather than independent feature-specific changes. Moving beyond the traditional focus on the spectral domain that has long dominated electrophysiological research^8,9,16^, this convergence highlights how diverse complexity, non-linear^11,13^, and spectral metrics provide complementary lenses for distinct biological processes that follow similar lifespan trajectories.

Spatially, age-related EEG dynamics were expressed along a continuum ranging from globally coordinated scalp-wide patterns to highly localized and spatially heterogeneous architectures. While some EEG descriptors exhibited remarkably consistent lifespan trajectories across nearly the entire scalp, others showed age-related effects confined to specific regions or even distinct trajectory classes across different scalp locations. The latter observation is particularly informative, as it demonstrates that the lifespan behavior of the same EEG feature can vary according to where it is measured. Such spatial diversity is consistent with evidence that brain maturation and aging proceed asynchronously across cortical systems, with structural, functional, and connectivity changes following region-specific developmental and aging trajectories rather than a single spatial program^28–30^.

These findings also have important implications for how lifespan EEG should be spatially represented. Global averaging provides an appropriate summary for descriptors whose age-related trajectories are coordinated across the scalp, as the averaged signal preserves their dominant lifespan profile–implying that high-density coverage or high-resolution arrays may be functionally redundant for tracking these specific features. In contrast, for descriptors exhibiting heterogeneous or spatially localized organizations, averaging can obscure regional trajectories, attenuate regionally pronounced age-related effects, or merge distinct developmental profiles into a single global estimate. The question is therefore not whether scalp-averaged or channel-level analyses are universally superior, but which spatial scale best reflects the pattern of the EEG descriptor under investigation.

Beyond trajectory shape, transition timing provides an additional dimension for understanding lifespan EEG. Many of the transition ages identified here occurred during early adulthood, indicating that substantial electrophysiological reorganization continues beyond adolescence. This timing is consistent with converging evidence that maturation of the human brain extends well into the third decade of life, including continued refinement of association cortices^29^, prolonged myelination and white-matter development^31^, and the maturation of executive cognitive systems supported by the prefrontal cortex^32^. Transition timing also differed between males and females, despite otherwise subtle sex-related differences. Males generally reached stabilization earlier, whereas females exhibited a later but more temporally concentrated transition window. This pattern is consistent with evidence that sex differences in brain development arise partly from differences in the timing and pace of maturation, influenced in part by pubertal neuroendocrine processes^33–36^.

By expressing each EEG descriptor relative to its own developmental timeline, we found that many distinct features followed similar regional sequences of transition across the scalp. Spectral, morphological, complexity, and nonlinear measures often reached their transition points in a comparable spatial order. This convergence suggests that transition timing is not specific to individual features, but may reflect broader neurodevelopmental processes that shape regional EEG maturation. This aligns with evidence that cortical development follows conserved spatial gradients and hierarchical principles, and with neuroimaging studies showing coordinated regional maturation across diverse brain phenotypes^3,28–30,37,38^.

The organizational framework proposed here changes how EEG alterations may be interpreted in studies of brain health and disease. Rather than asking only whether EEG features differ between healthy and clinical populations, this framework suggests asking whether the underlying organizational principles are altered. Pathological processes may preserve the overall trajectory while shifting transition ages, altering spatial organization, or disrupting the chronology of age-related change. Such alterations may remain undetected by conventional comparisons of feature magnitude alone, but become apparent when electrophysiological aging is considered across its temporal, spatial, and chronological organization.

Although this framework provides a broad population-level map of lifespan electrophysiological organization, several limitations should be considered. First, the cross-sectional design does not directly capture within-individual change and may be influenced by cohort effects; longitudinal studies are therefore needed to validate the inferred lifespan trajectories. Second, because the data were aggregated from independent cohorts, information on non-neurological medical conditions and medication use was not consistently available. This is particularly relevant in older adults, in whom systemic conditions, treatments, and preclinical neurodegenerative changes may influence EEG measures and complicate the definition of normative aging. FFuture studies should therefore use prospectively characterized cohorts and integrate EEG with clinical, cognitive, neuroimaging, demographic, and environmental factors. Replication in prospectively acquired, more tightly characterized cohorts will also be important. Finally, multivariate approaches could clarify how interactions among electrophysiological features shape lifespan brain organization.

In summary, this study shows that electrophysiological maturation and aging are organized through coordinated temporal and spatial patterns rather than simple linear shifts. By mapping how, where, and when EEG features change across the lifespan, we establish a normative framework for understanding the organization of human brain electrophysiology from childhood to late adulthood.

## Methods

### Dataset

Resting-state eyes-closed EEG data from 1,763 healthy participants (mean age = 40.1 ± 20.65 years, 54.4% female) were aggregated from 17 international datasets — comprising both open-access repositories and datasets obtained through collaborative agreements — covering a continuous age range from 5 to 85 years (Supplementary Materials Fig. S7). Detailed dataset references, access links, subjects’ description, and ethical approvals are provided in the Supplementary Materials.

### Preprocessing

Preprocessing followed the automated pipelines detailed in Ebadi et al.(2025) and Tabbal et al.(2025). EEG signals were first bandpass filtered (1-100 Hz) and down-sampled to 200 Hz^39,40^. Bad EEG channels were detected using the RANSAC method via *pyprep* algorithm^41^. Identified channels were subsequently corrected by spherical spline interpolation from neighboring electrodes. Signals were re-referenced to a common average reference. Eye-blink artifacts detection and removal was applied using Independent Component Analysis (ICA), and the IClabel algorithm^42^. A second bandpass filter (1–45 Hz) was applied before segmentation into 10-second epochs. Automated epoch rejection was performed using the Autoreject toolbox^43^, utilizing 10-fold cross-validation by default, with the exception of the ZenodoFuglsang^44^ dataset which utilized 5-fold cross-validation. Participants with a channel interpolation rate exceeding 20% were excluded from further analysis. This pipeline was applied uniformly across all datasets, except for the *Basel* cohort, which was preprocessed according to Yassine et al.^45^.

To address inter-site heterogeneity in channel configurations, all recordings were standardized to the 19-channel 10–20 montage prior to feature extraction, matching the minimum number of channels present across the contributing datasets.

### Feature Extraction

We extracted 103 scalp-level EEG features spanning the spectral, complexity, and morphological signal domains. Each feature was computed independently at each of the 19 channels of the standard 10-20 montage. In addition, a global estimate was obtained by averaging the feature values across all electrodes. This resulted in 20 spatial representations per feature (19 channel-specific and one global), yielding a total of 2,060 feature–channel combinations. Where applicable, features were computed for both broadband and across the five canonical frequency bands: delta (δ, 1–4 Hz), theta (θ, 4–8 Hz), alpha (α, 8–13 Hz), beta (β, 13–30 Hz), and gamma (γ, 30–45 Hz). To extend beyond conventional EEG features, we incorporated features from the catch22 toolbox — a filtered version of the Highly Comparative Time-Series Analysis (HCTSA) library^22^ — to capture complementary properties of EEG time series.

The spectral domain features included absolute and relative power across all five frequency bands, aperiodic components (intercept and slope), adjusted power (i.e., aperiodic-adjusted band power), alpha peak frequency, inter-band power ratios (θ/β, θ/α, δ/β, δ/α). It also included two frequency-related descriptors from the catch22 toolbox^22^—Centroid Frequency and Low-Frequency Power. The complexity domain comprised non-linear entropy measures, fractal dimensions computed across both broadband and canonical frequency bands, and autocorrelation-based properties, supplemented by catch22^22^ features quantifying incremental differences, signal periodicity, and forecasting error. The morphological domain captured the statistical shape of the EEG signal through envelope mean and standard deviation, and signals skewness and kurtosis — extracted for both broadband and canonical frequency bands — alongside Hjorth parameters and catch22^22^ distributional features quantifying the occurrence and magnitude of extreme signal events. The complete feature list is provided in Supplementary Materials Table S1.

Following feature extraction, we applied the neuroHarmonize framework^46^ to mitigate inter-site batch effects inherent to this multi-site dataset. This approach extends the standard ComBat algorithm^47^ by incorporating Generalized Additive Models (GAMs), enabling the removal of site-specific variance while preserving non-linear biological trajectories. Features were harmonized with *sites* modeled as the batch variable, while *age* and *sex* were included as protected covariates. The *age* covariate was treated as a non-linear smoothing term to accommodate the complex, multiphasic maturation and senescence patterns expected across the 5–85 year range.

### Multi-Model Inference

To identify the fundamental age-related trajectory for each feature, we utilized an Akaike Information Criterion (AIC) based Multi-Model Inference framework^19^. Each standardized EEG feature (Y, Z-scored to zero mean and unit variance) was fitted to four candidate models:

i. Null Model (Invariant): *Y* *= β*o + _β_s_S_ ɛ
ii. Linear Model: *Y* = *β*_o_ + *β*_1_. *Age* + *β_s_S* + ɛ
iii. Inverse Model: 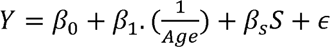
iv. Quadratic Model: *Y* = *β*_o_ + *β*_1_. *Age* + *β*_2_ . *Age*^2^ + *β_s_S* + ɛ

The inverse model captures rapid non-linear transitions followed by asymptotic stabilization. The quadratic model captures non-monotonic, symmetric trajectories with a single reversal point. The linear model served as a parsimonious baseline, and the null model represented the absence of any age-related change.

In the primary shape-selection phase, sex (*S*) was included as an additive categorical covariate in all candidate models to control for baseline differences while isolating the effect of age.

Model selection was governed by AIC, which identifies the most statistically parsimonious model within the candidate set by evaluating goodness-of-fit (*̂L*) while penalizing for model complexity (i.e., penalizing for the number of estimated parameters *(K)*^19^ :

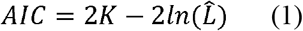

The “winning” model for each feature-channel combination was identified as the one yielding the minimum AIC value.

The AIC difference (ΔAIC) relative to the best-performing model was then computed for each candidate to quantify the relative evidence against each alternative. ΔAIC values were transformed into Akaike weights (*w_i_*) following Burnham and Anderson (2002)^19,48^. Akaike weights represent the probability that a given model is the best-approximating model within the candidate set, derived from the relative likelihood of each candidate:

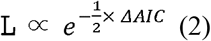

Weights were computed by normalizing these likelihoods across the four candidates, where i indexes the feature-channel combination:

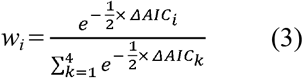

Analysis was restricted to feature-channel combinations reaching a selection certainty of *w_i_* ≥ 0.95 — mathematically equivalent to ΔAIC ≥ 6 — at which the winning model is at least 20 times more probable than any alternative.

### Normalized Effect Size

For each feature-channel combination, the magnitude of the age-related effect was quantified by the regression coefficient of the age term (β) from the winning model. Since all features were Z-scored prior to model fitting, β represents the change in standard deviation units per unit change in age, enabling direct comparison of effect magnitudes across features with heterogeneous scales and units. To further enable comparison across trajectory types — whose β parameters operate on different scales and are not directly commensurable across classes, we calculated the **Normalized Effect Size** of each feature by normalizing its coefficient against the maximum absolute β observed within its respective trajectory class (Linear, Quadratic, or Inverse):

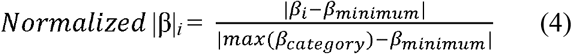

This transformation maps each feature’s effect magnitude to a normalized relative scale (0 to 1), where a value of 1 identifies the most influential feature within its functional aging category. Normalized effect sizes were categorized as low (|β|<0.20), medium (0.20<=|β|< 0.5), and high (|β|>=0.5)^49^. To preserve the directionality of age-related change, we computed the **Signed Normalized Effect Size** by incorporating the sign of the original coefficient:

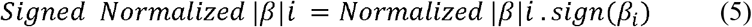

Positive and negative values indicate the direction of age-related change relative to the population mean. This allows the full landscape of aging effects—their relative magnitude, directionality, and functional trajectory class—to be visualized and compared systematically across the entire feature space, regardless of the underlying mathematical model.

### Evaluation of sex-related effects

Following identification of the optimal age trajectory for each feature, sex-related effects were evaluated by extending the winning trajectory model in two ways. First, an additive model assessed stable sex-dependent offsets across the lifespan while preserving the age trajectory. Second, an interaction model incorporated age-by-sex interaction terms (e.g., *Age* * *S* or 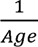 * *S*, depending on the winning trajectory) to determine whether males and females followed different developmental trajectories. Statistical significance of additive and interaction terms was assessed using Benjamini–Hochberg false discovery rate correction (FDR) (*q<0.05)*.

To complement significance testing with an estimate of biological relevance, effect sizes were quantified using Cohen’s f^2^, which quantifies the unique proportion of variance explained by either the additive sex term or the sex-age interaction, after accounting for the main effect of age constant:

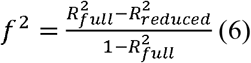

where 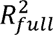 and 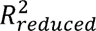 denote the coefficients of determination of the extended and base trajectory models, respectively. This metric provides a normalized measure of the magnitude of sex-related effects across features with heterogeneous lifespan trajectories^20^.

### Identification of Transition Points

To determine the precise timing of neurophysiological transitions, we performed a transition analysis on all EEG feature-channel combinations meeting the high model certainty threshold (*w_i_*≥ 0.95).

For features following a quadratic trajectory (y∼age + age^2^ ), the transition point was defined as the reversal point — the age at which the trajectory changes direction, computed as the vertex of the fitted parabola where the first derivative of the predicted trajectory (averaged across sexes) equals zero:

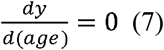

For features following an Inverse trajectory (*y*∼1/*age*), the milestone was defined as the age of stabilization — the age at which the rate of change became negligible (i.e., the feature entered a developmental plateau), identified as the point where the absolute first derivative fell below the standard error of the regression^50^:

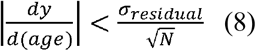

Where σ_residual_ is the standard deviation of the model residuals and *N* is the sample size.

Differences between male and female transition age distributions were quantitatively assessed using a two-sample K-S Test.

### Agreement Between Local and Scalp-Averaged Effects

To evaluate whether scalp-averaged EEG measures provide a valid proxy for localized lifespan dynamics, we performed a Bland–Altman analysis^21^ to assess the absolute agreement between local and global effect estimates. For each electrophysiological feature, the largest channel-specific normalized effect size (|β|) was compared with the corresponding effect derived from the scalp-averaged signal. The paired difference between these two estimates was plotted against their mean to quantify the systematic bias and the 95% limits of agreement (mean difference ± 1.96 × SD). This analysis determined whether the scalp-averaged signal systematically over- or underestimated the strongest localized age-related effect, thereby assessing its suitability as a global summary measure.

### Cross-channel synchronization of transition ages

Channel-specific transition ages were compared separately for each feature to assess how synchronized electrophysiological reorganization was across the scalp. For features represented at two or more channels, we calculated the difference between the earliest and latest transition ages as a measure of the total temporal span of scalp-wide reorganization. Median transition spans were then summarized across all features and within each spatial organizational mode.

To assess more typical regional synchronization, we also calculated the absolute difference in transition age for every pair of channels within each feature. These pairwise differences formed a distribution of temporal offsets. The interquartile range of this distribution was used to identify channel pairs with nearly synchronous transitions, which were then mapped to characterize the spatial organization of coordinated electrophysiological change.

### Spatiotemporal Clustering of Transition Profiles

To determine whether electrophysiological features with distinct mathematical definitions exhibit similar spatial patterns of developmental timing, we clustered features according to their regional transition profiles. First, transition ages were extracted for all robust feature–channel combinations (see “Identification of Transition Points”) and normalized across the 19 scalp electrodes for each feature. Pairwise Pearson correlations were then computed between the normalized spatial transition profiles of all features. To ensure reliable comparisons, correlations were calculated only for feature pairs sharing at least 11 significant electrodes (>50% of the standard montage). The resulting correlations were converted into a distance matrix (1 − r). Pairwise correlations that did not reach statistical significance (p ≥ 0.05) were assigned the maximum distance to prevent the clustering algorithm from grouping features based on weak or unreliable spatial similarity. Hierarchical agglomerative clustering with average linkage was subsequently applied to the resulting distance matrix.

## Data Availability

Details on dataset access, including repository links and any applicable access requirements, are provided in Supplementary Materials.

## Code Availability

The code used for the analyses in this study is publicly available on GitHub at: https://github.com/MINDIG-1/EEG-Lifespan-Trajectories

## Supplementary Materials

## Acknowledgements

The contributions of A.E., S.A. and M.H. to this work were supported by MINDIG as part of its research and development activities. The authors gratefully acknowledge the researchers and institutions who made their datasets available for research, as well as all participants who consented to the use of their data.

## Author contributions

M.K. performed the formal analysis and wrote the original draft of the manuscript. S.A., A.E., F.B., and M.H. supervised the work and contributed to reviewing and editing the manuscript. B.G., L.H., G.Y., I.K., V.P., P.G., B.R.Á., and M.V. contributed the datasets. U.G. and P.F. contributed the datasets and reviewed the manuscript. M.B. and M.D. provided administrative supervision. All authors approved the final version of the manuscript.

