## Supplementary Materials for "The Lifespan Architecture of Human EEG"

**Methods:**

**Datasets:**

Our cohort consisted of 1763 healthy controls. The data were aggregated from seventeen distinct studies, each detailed subsequently in the following sheet: [Dataset_Information](https://docs.google.com/spreadsheets/d/1oT7UR13NhGU2OsjKIxbHKfnZUWfGt5REySEhCRU_jGA/edit?usp=drive_link)

**Features:**

**Table. S1 |** List of Complexity, Spectral and Morphological features considered in this study

| **Complexity** | **Spectral** | **Morphological** |
| --- | --- | --- |
| Approximate Entropy | Adjusted Power-Alpha | Envelope Mean-Alpha |
| CATCH22 Embedding Distance | Adjusted Power-Beta | Envelope Mean-Beta |
| CATCH22 AC Function_timescale | Adjusted Power-Delta | Envelope Mean-Broadband |
| CATCH22 AC Function_First Minimum | Adjusted Power-Gamma | Envelope Mean-Delta |
| CATCH22 Mutual Information_Bins | Adjusted Power-Theta | Envelope Mean-Gamma |
| CATCH22 Time Reversibility | Aperiodic Intercept | Envelope Mean-Theta |
| Detrended Fluctuation-Alpha | Aperiodic Slope | Envelope Standard Deviation-Alpha |
| Detrended Fluctuation-Beta | Delta Alpha Ratio | Envelope Standard Deviation-Beta |
| Detrended Fluctuation-Broadband | Delta Beta Ratio | Envelope Standard Deviation-Broadband |
| Detrended Fluctuation-Delta | Peak Alpha Frequency | Envelope Standard Deviation-Delta |
| Detrended Fluctuation-Gamma | Power Absolute-Alpha | Envelope Standard Deviation-Gamma |
| Detrended Fluctuation-Theta | Power Absolute-Beta | Envelope Standard Deviation-Theta |
| CATCH22 Whiten Timescale | Power Absolute-Delta | Hjorth Activity |
| CATCH22 Forecast Error | Power Absolute-Gamma | Hjorth Complexity |
| Higuchi Fractal-Alpha | Power Absolute-Theta | Hjorth Mobility |
| Higuchi Fractal-Beta | Power Relative-Delta | Kurtosis-Alpha |
| Higuchi Fractal-Broadband | Power Relative-Alpha | Kurtosis-Beta |
| Higuchi Fractal-Delta | Power Relative-Beta | Kurtosis-Broadband |
| Higuchi Fractal-Gamma | Power Relative-Gamma | Kurtosis-Delta |
| Higuchi Fractal-Theta | Power Relative-Theta | Kurtosis-Gamma |
| Hurst Exponent C | Total Power-Alpha | Kurtosis-Theta |
| Hurst Exponent H | Total Power-Beta | Skewness-Alpha |
| CATCH22 Automutual information | Total Power-Broadband | Skewness-Beta |
| Katz Fractal-Theta | Total Power-Delta | Skewness-Broadband |
| Katz Fractal-Alpha | Total Power-Gamma | Skewness-Delta |
| Katz Fractal-Beta | Total Power-Theta | Skewness-Gamma |
| Katz Fractal-Broadband | CATCH22 Low Frequency Power | Skewness-Theta |
| Katz Fractal-Delta | CATCH22 Centroid Frequency | CATCH22 Distribution_Outlier_positive |
| Katz Fractal-Gamma | Theta Alpha Ratio | CATCH22 Distribution_Outlier_negative |
| Lempel-Ziv | Theta Beta Ratio | CATCH22 Distribution_Histogram_mode 10 |
| CATCH22 High Fluctuation |  | CATCH22 Distribution_Histogram_mode 5 |
| CATCH22 Periodicity |  | CATCH22 Distribution_Mean |
| Sample Entropy |  | CATCH22 Distribution_Standard Deviation |
| Spectral Entropy |  |  |
| CATCH22 Stretch Decreasing |  |  |
| CATCH22 Stretch High |  |  |
| CATCH22 Entropy Pairs |  |  |
| CATCH22 Transition Variance |  |  |
| CATCH22 dfa |  |  |
| CATCH22 Rescaled Range Analysis |  |  |

**Complexity Features**

*Descriptors quantifying signal irregularity, temporal dependence, fractal organization, predictability, and nonlinear dynamics.*

**Entropy-based complexity:** Approximate Entropy, Sample Entropy, Spectral Entropy

Capturing: signal irregularity, predictability and information richness.

**Fractal scaling & long-range temporal correlations:** Detrended Fluctuation Analysis (Alpha, Beta, Delta, Gamma, Theta, Broadband), Hurst Exponent (C), Hurst Exponent (H)

Capturing: long-range temporal dependence and self-similar neural dynamics.

**Fractal geometry:** Higuchi Fractal Dimension, Katz Fractal Dimension (all frequency bands)

Capturing: geometric complexity of EEG waveforms.

**Algorithmic complexity:** Lempel–Ziv Complexity

Capturing: sequence compressibility and information complexity.

**Catch22 nonlinear dynamics,** all details are presented by Lubba et al., 2019. **[23]**

**Spectral Features**

*Descriptors characterizing oscillatory activity, spectral composition, and background aperiodic activity.*

**Absolute Power, Relative Power, Adjusted Power:** Delta, Theta, Alpha, Beta, Gamma

**Total Power:** Delta, Theta, Alpha, Beta, Gamma, Broadband

**Oscillatory ratios:** Delta/Alpha, Delta/Beta, Theta/Alpha, Theta/Beta

**Oscillation frequency:** Peak Alpha Frequency

**Aperiodic activity:** Aperiodic Intercept, Aperiodic Slope

**Morphological Features**

*Descriptors quantifying statistical properties and amplitude fluctuations of EEG waveforms.*

**Hjorth descriptors:** Hjorth Activity, Hjorth Mobility, Hjorth Complexity

Capturing: signal variance, waveform characteristics.

**Envelope statistics: Mean & Standard deviation:** Delta, Theta, Alpha, Beta, Gamma, Broadband

Capturing: average oscillatory amplitude and amplitude variability.

**Distribution shape:**

**Skewness & Kurtosis:** Delta, Theta, Alpha, Beta, Gamma, Broadband

Capturing: asymmetry and peak of EEG amplitude distributions.

**Spatial and Topographical Organization**

To investigate the spatial heterogeneity of aging, EEG metrics were analyzed at three distinct spatial scales: global, lobe, and local (electrode-level). The global scale was defined as the arithmetic feature-wise mean across the 19 electrodes. For the lobe scale, EEG features across the electrodes were aggregated into five functional regions: **frontal** (Fp1, Fp2, F3, F4, F7, F8, Fz), **central** (C3, C4, Cz), **parietal** (P3, P4, Pz), **temporal** (T3, T4, T5, T6), and **occipital** (O1, O2). This hierarchical approach allowed us to identify whether non-linear trajectories were brain-wide phenomena or localized to specific functional regions.

**Results:**

[Supplementary Table 1](https://docs.google.com/spreadsheets/d/1aPxy2Dx5o7bZ605EFXunCmnPcO3vXm9p69SJ4iZi-sw/edit?usp=drive_link)

[Supplementary Table 2](https://docs.google.com/spreadsheets/d/1vcxLvF1oXvKmV2y5Q8OLp3EJRsv7qO19pOmHJF4pVVI/edit?usp=drive_link)

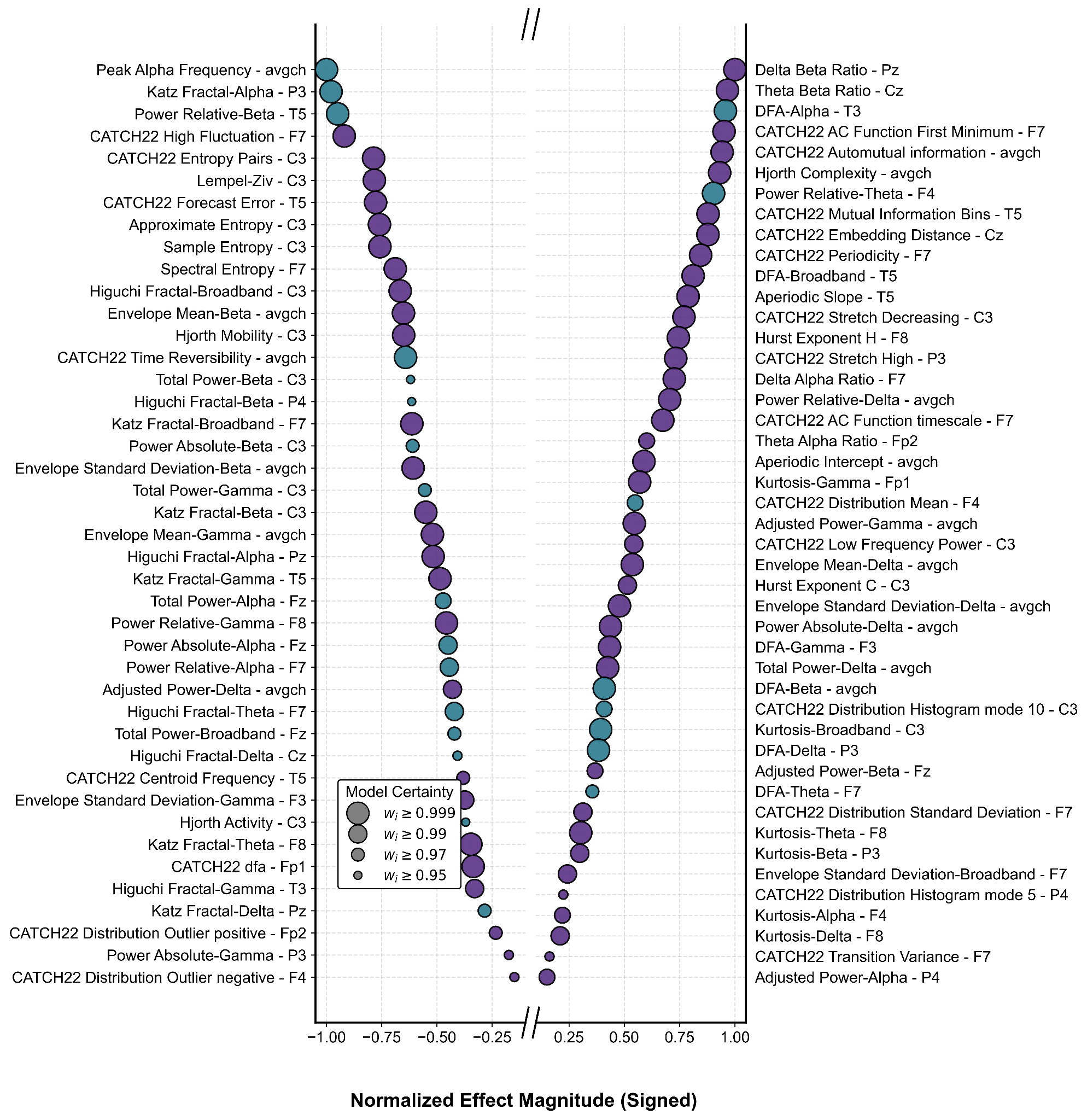

**Fig. S1 | Extended Version: Signed normalized effect size for the most significant channel-specific features.** The horizontal axis represents normalized effect sizes ranging [-1.0 1.0]. Data point size is proportional to model certainty, while color indicates the winning trajectory model (Inverse, purple; Quadratic, teal).

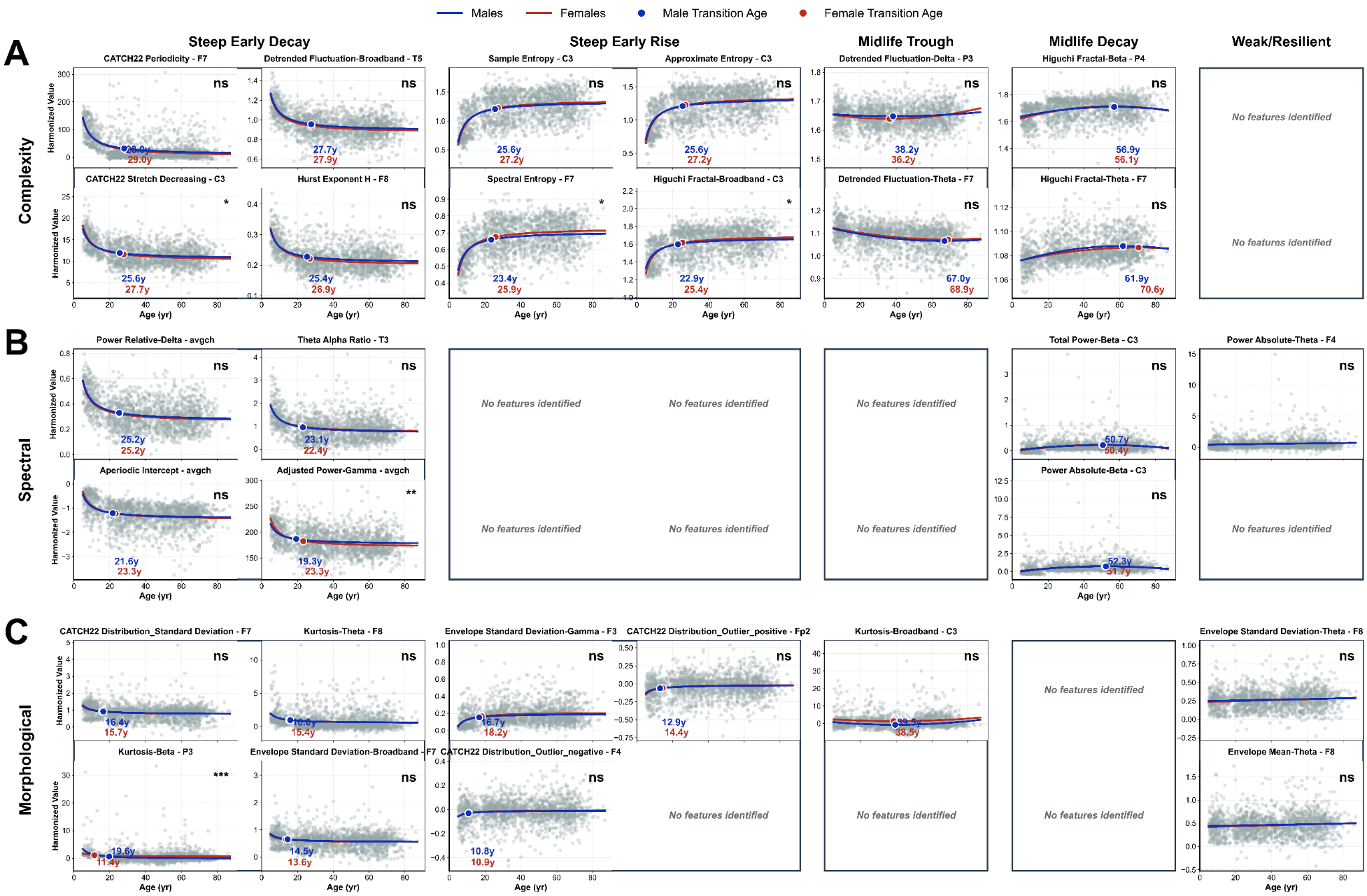

**Figure S2 | Additional sex-stratified lifespan trajectories across electrophysiological feature domains.** Shown are additional electrophysiological descriptors selected immediately below those presented in Fig. 1 for each trajectory class within the complexity (A), spectral (B), and morphological (C) feature families. Blue and red curves represent fitted lifespan trajectories for males and females, respectively, with colored markers indicating the corresponding transition ages. Statistical annotations indicate sex differences in transition age (P < 0.05, P < 0.01; ns, not significant). Empty panels denote trajectory class–feature family combinations for which no additional descriptors were identified.

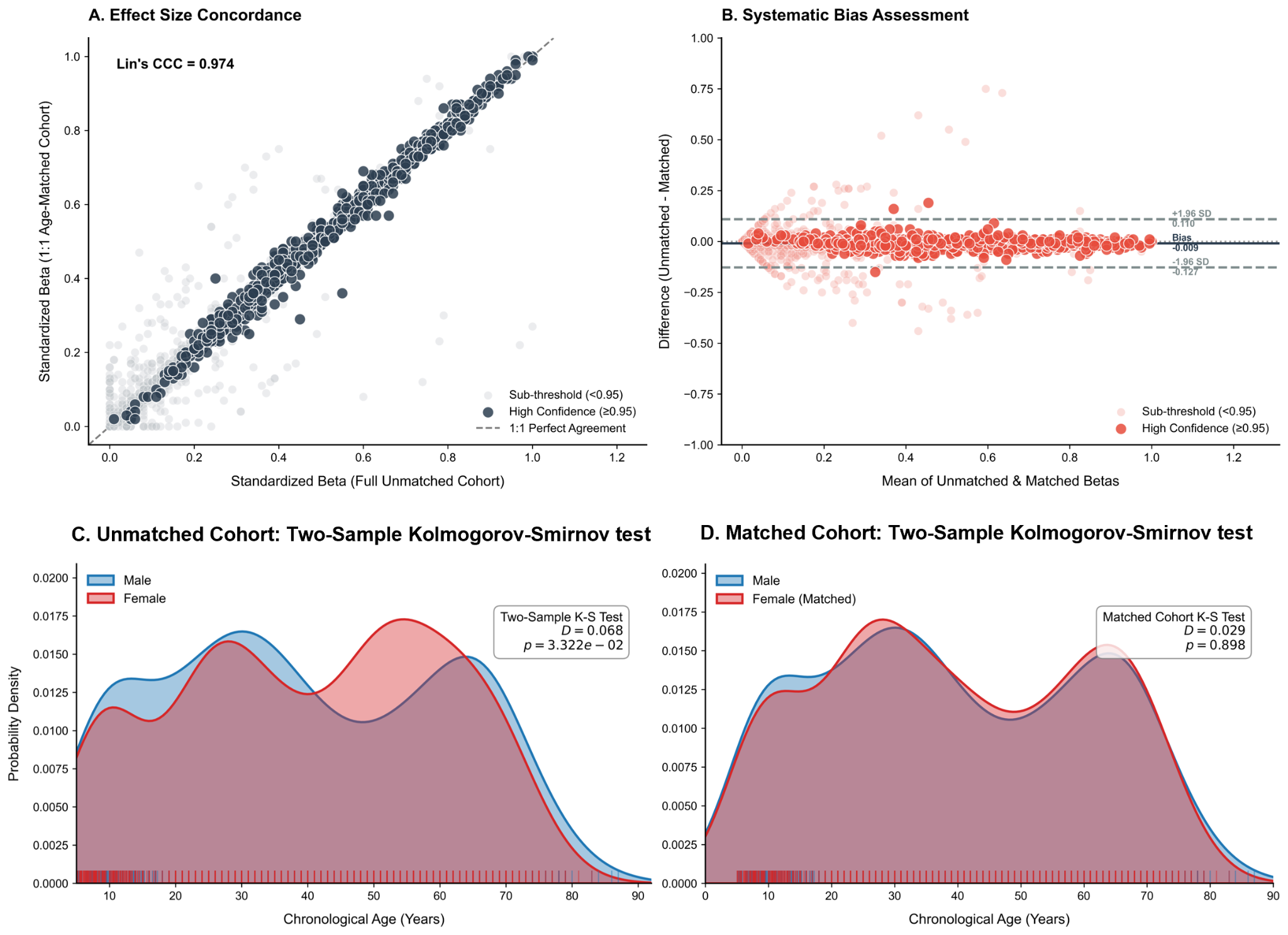

**Figure S3 | Validation of age-related trajectory robustness following sex-stratified age-bin matching.** (A) Concordance between standardized effect sizes (β) estimated from the full unmatched cohort and the age-matched cohort after matching participants within age bins stratified by sex. Each point represents one electrophysiological descriptor. Dark points indicate features with high model-selection certainty (Akaike weight ≥ 0.95), whereas light points denote sub-threshold features. The dashed diagonal represents perfect agreement (1:1), and Lin's concordance correlation coefficient (CCC) quantifies overall agreement between cohorts. (B) Bland–Altman analysis [21] comparing standardized effect sizes from the unmatched and matched cohorts. The solid horizontal line indicates the mean bias, while dashed lines denote the 95% limits of agreement (±1.96 SD), demonstrating minimal systematic bias after matching. (C) Kernel density estimates of the chronological age distributions for male (blue) and female (red) participants in the original cohort, showing a significant difference in age distributions between sexes (two-sample Kolmogorov–Smirnov test [53]). (D) Age distributions after sex-stratified age-bin matching, demonstrating successful balancing of the male and female age distributions (Kolmogorov–Smirnov test, P = 0.898). Collectively, these analyses confirm that the observed age-related electrophysiological trajectories are robust to demographic imbalance and are not driven by unequal age or sex distributions across the cohort.

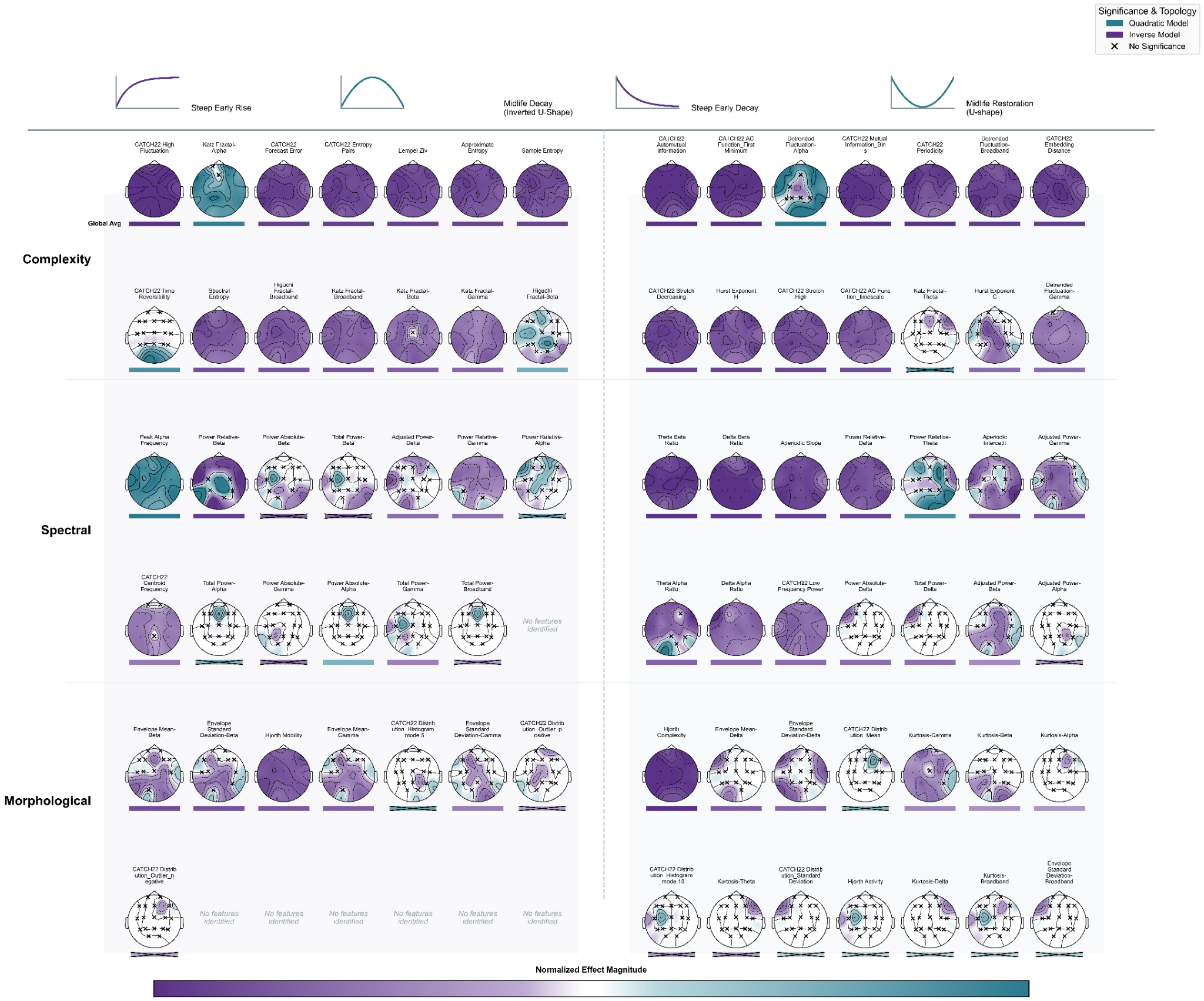

**Figure S4 | Extended multiscale topography of age-related changes across representative electrophysiological features.** Extended version of Fig. 3A, showing additional representative electrophysiological features exhibiting high model certainty (wᵢ ≥ 0.95). Features are grouped by domain (complexity, spectral, and morphological) and organized according to their dominant lifespan trajectory class: steep early rise, midlife restoration (U-shape), steep early decay, and midlife decline (inverted U-shape). Each topographic map depicts the regional distribution of normalized effect magnitudes, while the horizontal bar beneath each map summarizes the corresponding scalp-averaged effect. Channels not meeting the model-certainty threshold are indicated by × symbols. This figure complements Fig. 3A by illustrating additional representative examples spanning the three electrophysiological feature domains.

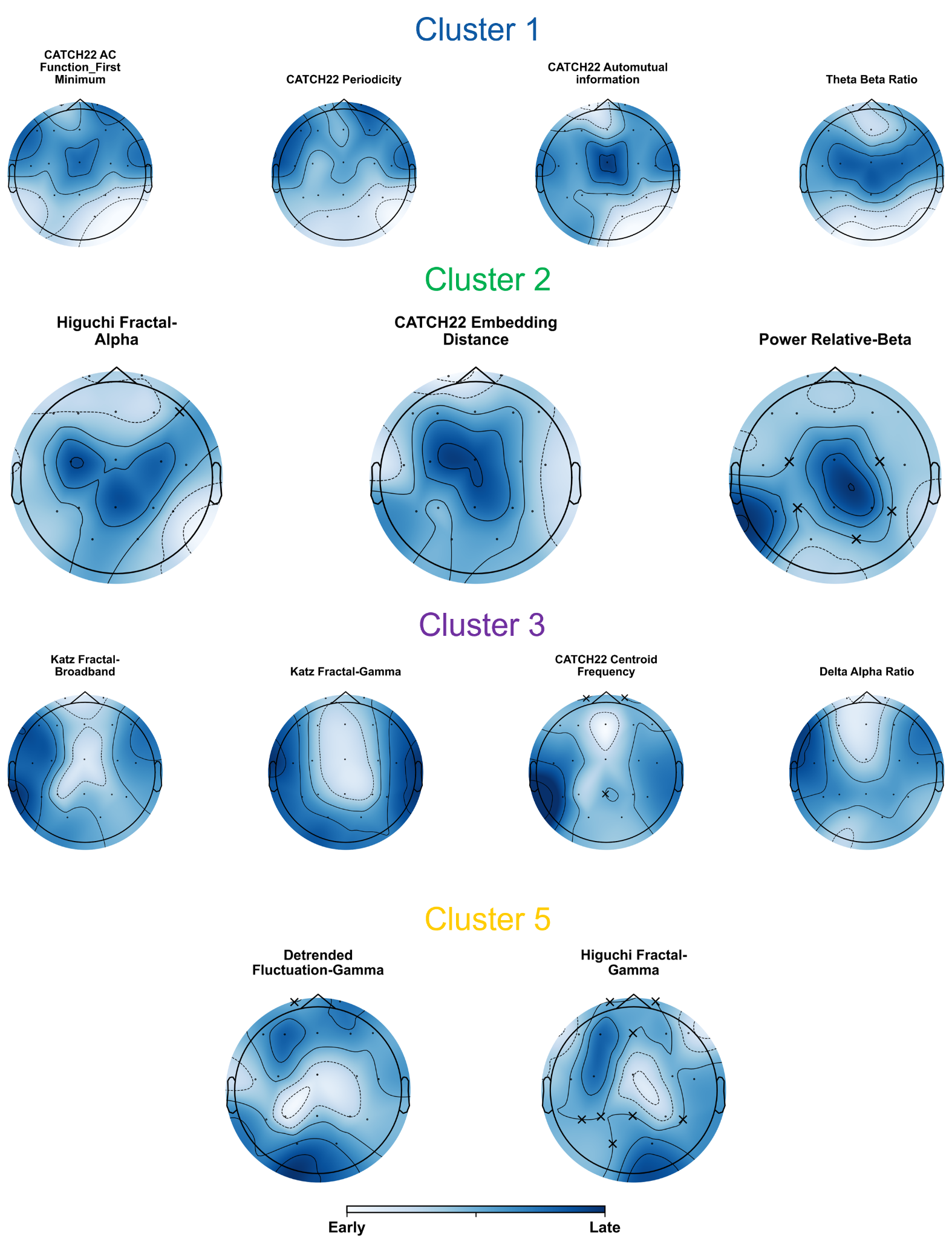

**Supplementary Figure S5 | Representative transition-age topographies for the conserved chronological transition clusters.** Representative normalized transition-age maps for electrophysiological features assigned to **Clusters 1, 2, 3 and 5**, illustrating the characteristic spatial organization of transition timing within each cluster. Features belonging to the same cluster exhibit similar regional sequences of early-to-late electrophysiological transitions despite differing in their mathematical definition. Crosses (×) denote channels not meeting the model-certainty threshold (*w*ᵢ < 0.95). Color intensity represents normalized transition timing, ranging from earlier (light) to later (dark) transitions.

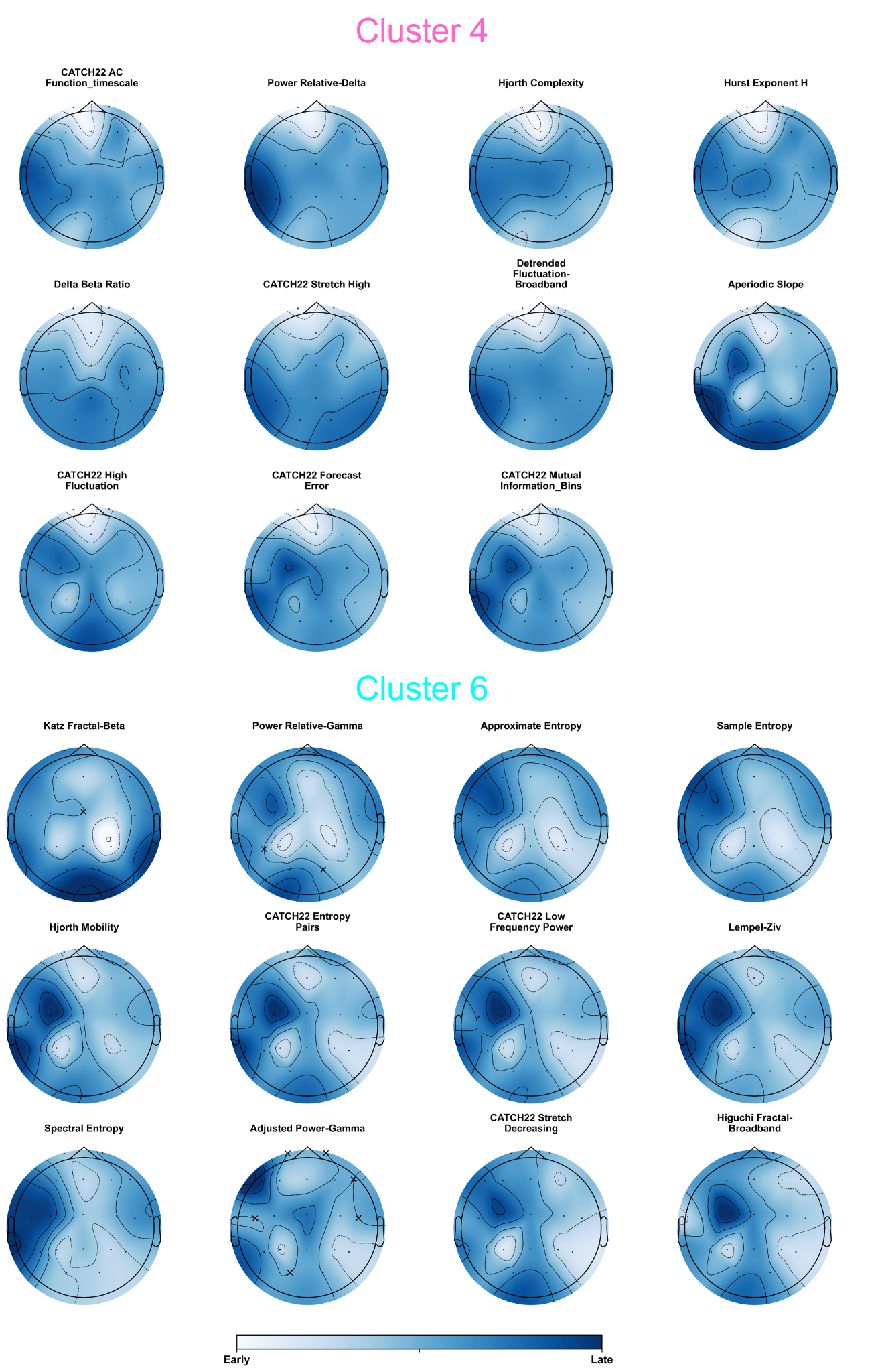

**Supplementary Figure S6 | Electrophysiological features comprising the 4th and 6th clusters.**

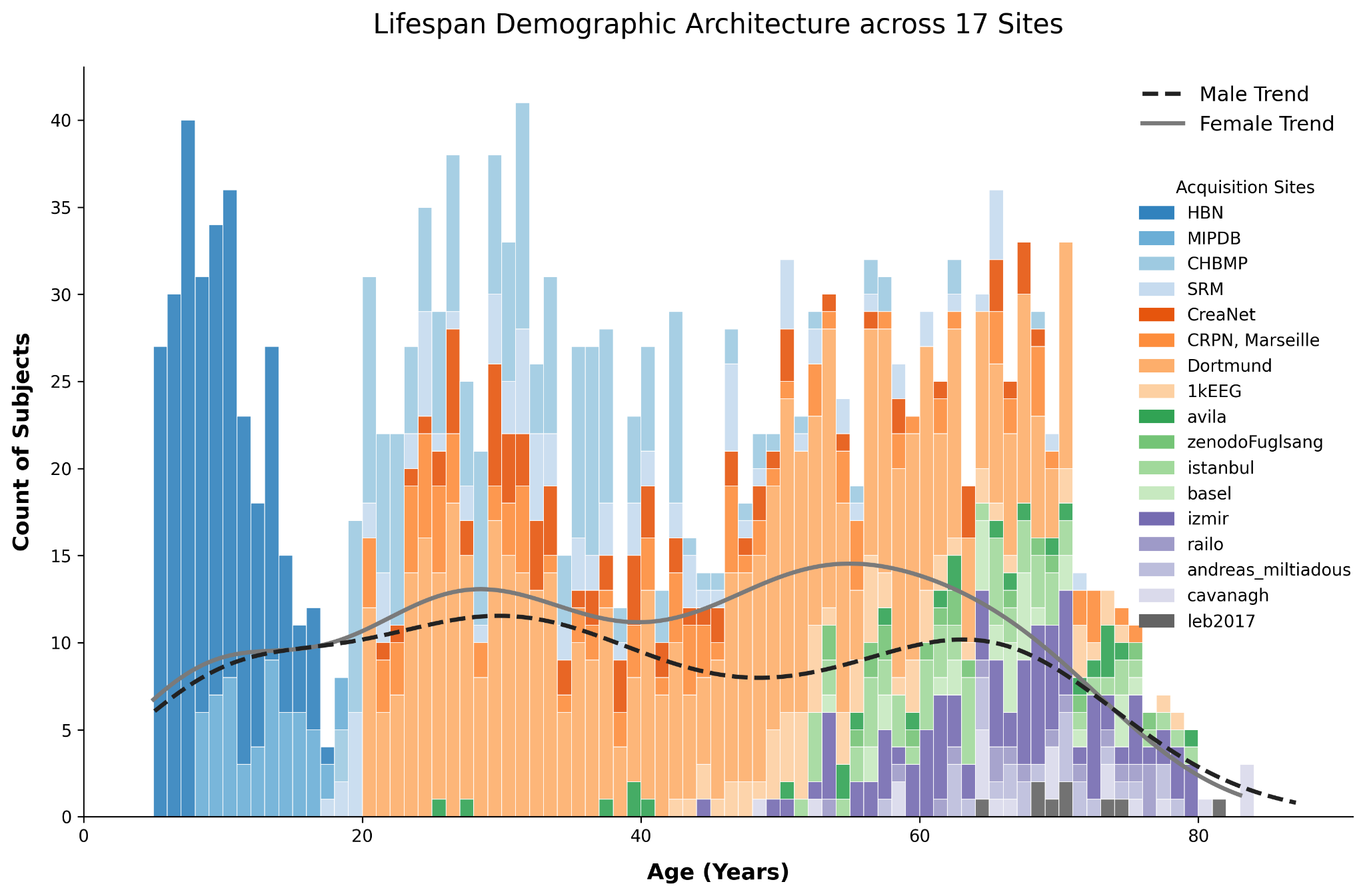
**Supplementary Figure S7 | Demographic composition of the harmonized lifespan EEG cohort.** Stacked histogram showing the age distribution of 1,763 healthy participants (5–85 years) included in the study across 17 independent acquisition sites. Superimposed curves represent kernel density estimates of the age distributions for male (black dashed line) and female (gray solid line) participants, demonstrating broad age coverage in both sexes.
